# A Public Database on Traumatic Brachial Plexus Injury

**DOI:** 10.1101/399824

**Authors:** Cristiane B Patroclo, Bia L Ramalho, Juliana S Maia, Maria Luíza Rangel, Fernanda F Torres, Lidiane Souza, Kelly R Braghetto, Claudia D Vargas

## Abstract

We hereby present the first worldwide public digital database centred on adult Traumatic Brachial Plexus Injury (TBPI). This initiative aims at reducing distance between clinical and experimental practice and encouraging data sharing and reuse. Detailed electronic questionnaires made with the free software LimeSurvey were designed to collect patients’ epidemiological, physical and clinical data. The freely available software Neuroscience Experiments System (NES) was employed to support data storage and management. First results of this effort concern data collected from 109 Brazilian adult TBPI patients with varying degrees of functional impairment. The sample is composed by large majority of men (84.4%), mean age of 32.1 (11.3 SD) years old, victims of motorcycle accidents (67%). The similarity of this dataset basic descriptors with those from previous reports in TBPI validates the strategies employed herein. Managing data from diverse provenance in TBPI may allow identifying functional markers related to the patients’ clinical improvement and foster the development of new investigative tools to unveil its mechanisms.

## 1 Introduction

Construction, maintenance and curation of public databases are becoming fundamental to propel our understanding of the nervous system function and dysfunction. This data-sharing paradigm emerged in the literature and media in the 90s^(1,2)^ with the International Consortium for Brain Mapping, the first major data sharing initiative for fMRI measures. The growth of this trend had only been possible by the technological progress that substantially increased the capacity to generate data in neuroscience and the huge development of information technology^(3,4)^. In accordance to the data-sharing paradigm, we devised a public database able to store data of diverse provenance in the domain of traumatic brachial plexus injury (TBPI) and its surgical reconstruction in adult patients.

The brachial plexus is a nerve net that congregates the ventral divisions from C5 to T1 spinal nerves and is responsible for ipsilateral upper limb motricity and sensibility^(5)^. TBPI mainly affects young males usually involved in motorcycle accidents, often leading to severe motor and sensitive impairment in the affected upper limb^(6)^. Psychic, social and quality of life impairments are also reported^(7,8)^.

TBPI’s treatment of choice is the surgical reconstruction allied with physical therapy^(9,10)^. Muscle strength is by far the focus of most frequent treatment outcomes^(11,12)^.

TBPI main associated prognostic factors are: patient’s age, lesion site, severity and mechanisms, associated traumatic lesions, time interval between the injury occurrence and surgery and the employed surgical repair technique^(13–23)^. However, the lesion complexity, its heterogeneity, and the well documented brain plasticity that follows a peripheral nerve injury^(24–26)^, make it very difficult to preview patients’ outcomes. This argues for the incorporation of other measures of success after TBPI^(12)^ and for the development of specific instruments for the functional evaluation of these patients^(27)^.

In this context, for the first time a set of detailed questionnaires designed to collect TBPI patients’ epidemiological, clinical, physical and surgical data together with the first set of anonymized results concerning 109 TBPI patients are made publicly available, as described here.

## 2 Methods

The development of the TBPI database was carried out by a multidisciplinary team comprising physicians, physiotherapists, neuroscientists and computer scientists.

Database building efforts involved the following steps: 2.1 patients selection, 2.2 data selection, 2.3 electronic questionnaires development, 2.4 data management platform development, 2.5 data entering, 2.6 de-identification of personal data, 2.7 data access.

### 2.1 Patient selection

All the TBPI patients included in the database were older than 18 years and were evaluated at the Institute of Neurology Deolindo Couto of the Federal University of Rio de Janeiro (INDC-UFRJ) from 2010 to 2017 by physicians and physical therapists. Patients prospectively evaluated voluntarily gave their written consent allowing the publication of their de-identified data in a public database. Retrospective data, collected by the same group before the TBPI Database project, was also allowed to be included in the database by the local Ethics Committee.

### 2.2 Data selection

Epidemiological, clinical, physical and surgical data were collected through specifically designed questionnaires named: Unified Admission Assessment, Unified Follow up Assessment and Unified Surgical Evaluation.

Unified Admission Assessment (UAA) concerns information about TBPI (side, causes and associated lesions), ongoing treatments (physiotherapy, orthosis, medication), neurological exam with visual inspection (glenohumeral subluxation, scoliosis, Horner’s syndrome, swelling, scares and trophic changes), presence of Tinel sign, sensory evaluation (light touch, pain, joint position sense, kinesthesia and pallesthesia), motor evaluation (motion range and muscle strength), pain occurrence and lesion site (preferably based on surgical information, followed by complementary exam, previous notes and physical exam). UAA was filled at the first interview and relied on medical records for missing data.

Unified Follow up Assessment (UFA) reviews ongoing treatments and neurological exam previously evaluated with UAA. It was filled at follow-up visits, ideally at each six months.

Unified Surgical Evaluation (USE) details surgical findings (level and type of lesion) and the employed procedures (type of surgery with its specificities). The USE questionnaire is filled by the neurosurgeons just after the surgery or by the researchers based on medical records.

### 2.3 Electronic questionnaires development

Electronic questionnaires present advantages when compared to paper-based ones such as ease and speed of administration, enforcement of data standardization (e.g., by using fixed choice response formats), immediate connection with the database, easier access to data and efficiency and security in data storage^(28)^. The questionnaires described in the previous section were created in an electronic format using the open-source survey system LimeSurvey. The decision for choosing LimeSurvey stemmed from the free availability of the tool and the fact that it relies on an underlying database management software, which can be deployed on a server that is deemed appropriate to store the target data and customized to support different data access policies. In addition, it supports several question types, enables the definition of restrictions on questions, admits design of logical branching based on answers and scores, allows the creation of multilingual surveys, enables the user to export collected data into spreadsheets, and supports general survey security settings.

The created electronic questionnaires follow the general structure of the paper-based questionnaires previously used by the group, facilitating adherence. The UAA, UFA and USE questionnaires favor multiple-choice questions to avoid excessive answers variability and enforce standardization in data entering. Besides, conditional branching arrangement of questions were employed to ensure data detailing.

### 2.4 Data management platform development

To meet the TBPI project’s demands for data acquisition, management and sharing, a software tool named Neuroscience Experiment System (NES) was developed. NES is a free, safe, user-friendly platform originally devoted to assist researchers in their experimental data collecting routine and to enable experiments reproduction. The platform keeps comprehensive and detailed descriptions of experimental protocols in a unified repository, as well as data and metadata from different provenance, including epidemiological, clinical and physical database described in the present report. With this aim, NES was integrated with LimeSurvey to facilitate questionnaires’ administration and to centralize data access.

### 2.5 Data entering

Prospective TBPI patients’ data were collected through personal interview conducted by a neurologist and/or a physiotherapist. The collected data was directly registered in the database through the NES platform. Retrospective TBPI patients’ data required careful selection, cleaning and transformation before entry the database. This process was done by the same neurologist and/or a physiotherapist who collected prospective data. Also, different sources were used in a hierarchical way to ensure data completeness and consistency: previous research records, medical records and patients’ report via phone call.

### 2.6 De-identification of personal data

Patients’ identification is only known by the local facility researchers. For the public version of the database the following information are suppressed: name; birthdate; ethnicity; address; phone number; professional information; dates (replaced by time intervals); injury circumstances as local of occurrence and details of hospitalization; and surgical and treatment places. Each patient in the database is identified by a noninformative code automatically generated by NES. These safety actions were inspired by the “Guidelines for working with small numbers”^(29)^ from the Washington State Department of Health and by Hrynaszkiewicz (2010)^(30)^.

### 2.7 Data access

The anonymized version of the TBPI database (exported from NES) is available to public access in an Open Database portal https://neuromatdb.numec.prp.usp.br/experiments/brachial-plexus-injury-database-v2/. Periodic database updates will be available in the portal with a brief description of the changes in the updated data as compared to the previous version.

The data are also hosted at figshare.

All public experimental data in the NeuroMat Open Database is available under the Creative Commons Attribution 4.0 International (CC BY 4.0) (https://creativecommons.org/licenses/by/4.0/) license. Therefore, to be able to download data, a potential user is required to agree and accept the CC BY 4.0 terms. According to CC BY 4.0, licensees may copy, distribute, and display the data and generate derivative works based on it only if they give the author(s) the appropriate credits (attribution).

## 3 Code availability

NES version 1.39 was used to create and manage the BPTI database. NES is a free software licensed under Mozilla Public License version 2.0. Its source code and documentation are available at https://github.com/neuromat/nes.

LimeSurvey version 2.05 was used to create and manage the electronic questionnaires. Its source code and documentation are available at http://www.limesurvey.org/. The structures of LimeSurvey questionnaires are stored in a .LSS file (which is basically an XML file) and can be imported in Limesurvey platform to (re)create a questionnaire.

## 4 Data Records

No specific software is required to handle the TBPI data from the NeuroMat Open Database. The dataset is downloaded as a single .ZIP file. It compresses several directories with plain-text files containing textual and numeric data (e.g., .CSV files for tabular data and .JSON files for metadata) that can be opened with a large variety of computational tools, from simple text editors to general-purpose statistical softwares.

In the root directory of the decompressed dataset, the data and metadata files are organized according the following hierarchical structure:

~~~
− Citation.txt
− Experiment.csv
− License.txt
+ Group_patients-with-brachial-plexus-injury (directory)
− Participants.csv
+ Experimental_protocol (directory)
   − Experimental_protocol_description.txt
   − Experimental_protocol_image.png
   + Step_1_questionnaire ≫ Additional_files (directory)
      − unified-admission-assessment.lss
      − unified-admission-assessment.pdf
   + Step_2.1_questionnaire ≫ Additional_files (directory)
      − unified-surgical-evaluation.lss
      − unified-surgical-evaluation.pdf
   + Step_2.2_questionnaire ≫ Additional_files (directory)
      − unified-follow-up-assessment.lss
      − unified-follow-up-assessment.pdf
+ Questionnaire_metadata (directory)
   + Q44071_unified-admission-assessment (directory)
      − Fields_Q44071_en.csv
      − Fields_Q44071_pt-BR.csv
   + Q61802_unified-surgical-evaluation (directory)
      − Fields_Q61802_en.csv
      − Fields_Q61802_pt-BR.csv
   + Q92510_unified-follow-up-assessment (directory)
      − Fields_Q92510_en.csv
      − Fields_Q92510_pt-BR.csv
+ Per_questionnaire_data (directory)
   + Q44071_unified-admission-assessment (directory)
      − Responses_Q44071.csv
   + Q61802_unified-surgical-evaluation (directory)
      − Responses_Q61802.csv
   + Q92510_unified-follow-up-assessment (directory)
      − Responses_Q92510.csv
+ Per_participant_data (directory)
   + Participant_PXXXX (directory)
      + Step_1_questionnaire (directory)
         − Q44071_unified-admission-assessment.csv
      + Step_2.1_questionnaire (directory)
         − Q61802_unified-surgical-evaluation.csv
      + Step_2.2_questionnaire (directory)
         − Q92510_unified-follow-up-assessment.csv
   + Participant_PYYYY (directory)
      (…)
    (…)
~~~

*Citation.txt* and *License.txt* contain information about the license under which the dataset has been published and how it must be cited.

*Experiment.csv* is a CSV (*comma-separated values*) file with basic information (name, description, start and end date) about the experiment and study which have generated the dataset. In all the .CSV files of the dataset, values (fields) in each line are separated by commas (,), textual values are enclosed by quotation marks (*“”*), and the first file line contains the column titles (i.e., the descriptions of fields).

The directory *Group_patients-with-brachial-plexus-injury* contains data and metadata related to the group of subjects called “Patients with brachial plexus injury” (which is the only group in the dataset) and its experimental protocol. The files *Participants.csv* contains personal, non-sensitive information about the subjects: participant code and gender. The *Experimental_protocol* directory contains a text file and an image that summarizes the procedure used to gather the data (i.e., the sequence of administration of the questionnaires). In addition, *Experimental_protocol* contains directories for each one of the three questionnaires used to collect the patient’s data (Step 1 - *Unified Admission Assessment*, Step 2.1 - *Unified Surgical Evaluation*, and Step 2.2 - *Unified Follow-up Assessment*). In each of these directories, the respective questionnaire can be found both in the .PDF and .LSS file formats. The .LSS file is from LimeSurvey; it can be used to recreate (import) the questionnaire structure in any LimeSurvey server.

In the *Questionnaire_metadata* subdirectory, one may found .CSV files which describe the complete structure of the three questionnaires both in English and Brazilian Portuguese. Each .CSV file contains, for each question in the questionnaire, its identifying code, description (complete text), type, sub-questions and answer options. To facilitate the understanding and manipulation of data collected using the questionnaires, we adopted a naming convention to question codes: the prefix of the code indicates the type of data collected by the question. For example, for the question “How old is the patient?”, *intAge* should be used as code. The “int” prefix indicates that the value for age is an integer number. Table 1 shows the data types of the questions which appear in our questionnaires and their respective code prefixes.

**Table 1.**
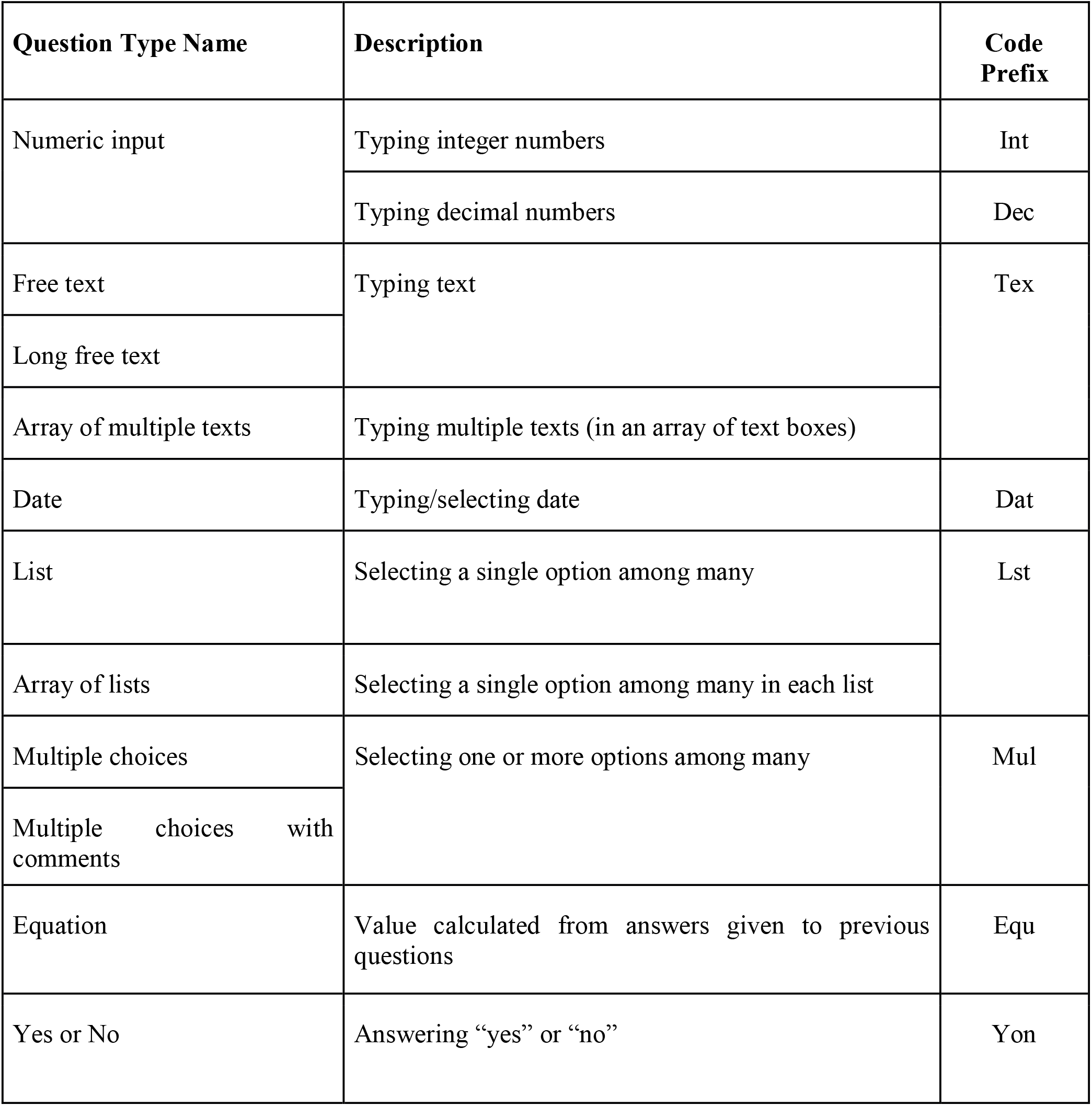
LimeSurvey question types and their respective prefix codes.

The patients’ data collected by means of the electronic questionnaires is replicated in two directories: *Per_questionnaire_data* and *Per_participant_data.* In the former, for each one of the three questionnaires, there is a directory containing a .CSV file (e.g., *Responses_Q44071.csv*) with all the patient responses collected through the questionnaire. Each line in these CSV files contains a response for a particular patient; the identification of the patient appears in the column entitled *“participant_code”*. Each patient has exactly one response for the *Unified Admission Assessment*, but he/she can have none or more responses for the *Unified Surgical Evaluation* and *Unified Follow-up Assessment*. In other words, the participant code is unique in the *Responses_Q44071.csv* file, but not in *Responses_Q61802.csv* and *Responses_Q92510.csv*. Different lines in the .CSV file with a same participant code indicate different responses of a same patient for a same questionnaire.

In the *Per_participant_data* directory, there is directory for each patient who participated in the study, containing a .CSV file for each questionnaire filled for him/her. If a patient has multiple responses for a given questionnaire, they will appear as different lines in its .CSV file. Each CSV file in the *Per_participant_data* directory contains responses of only one patient.

In the .CSV file containing responses for a given questionnaire, each column corresponds to a question or a sub-question of the questionnaire. All information required to understand the meaning of a column and its values can be recovered from the questionnaire’s metadata file through the column name. Searching by a response column name in the “question_index” column of the questionnaire’s metadata file, one will find one or more lines which describe the correspondent question or subquestion and the type of responses it accepts.

## 5 Technical Validation

To our knowledge, this is the first public dataset concerning adult TBPI, thus precluding its comparison with other databases. Therefore, in this section we will limit our data validation to the basic epidemiological descriptors available in literature.

The present dataset contains data from 109 Brazilian adult patients with TBPI. In agreement with the classical report of Narakas^(17)^, and subsequent national^(31–34)^ and international series(19,20,35), our sample is composed mainly by young (average age at injury: 32.1 years ± 11.3SD) male (84.4 %) subjects involved in traffic accidents. Fifty-six patients (51%) have left side injury. Figure 1A depicts results of TBPI causes in 114 nerve injuries from 109 patients (one patient had bilateral injury and four patients had unilateral injury provoked by two different causes). The entire brachial plexus (encompassing C5 to T1 roots) is affected in 44% of the patients. Four patients present more than one site of lesion (Figure 1B).

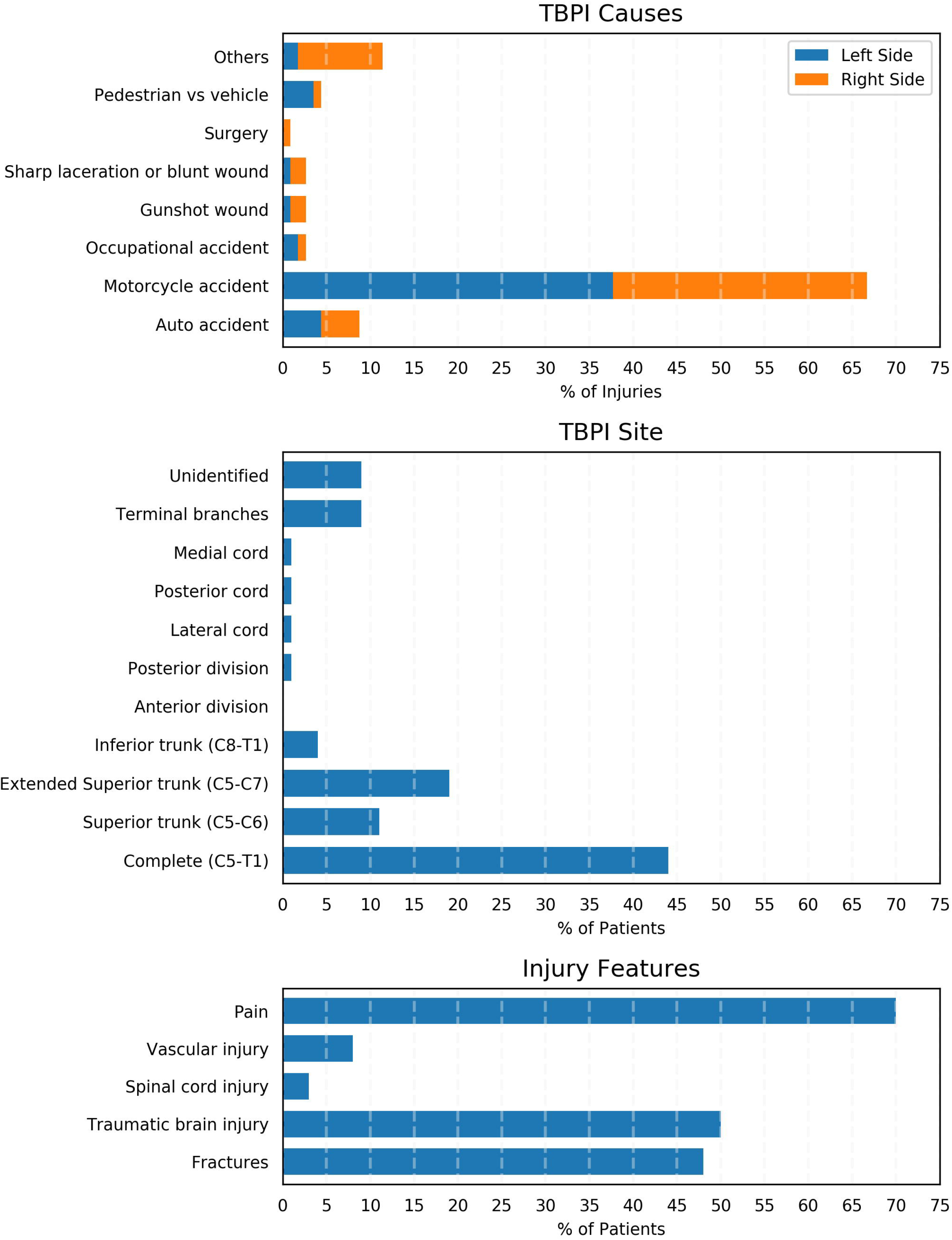

The percentage of associated injuries (fractures and traumatic brain, spinal cord and vascular injuries) and reported pain in the 109 patients (Figure 1C) are also in accordance with the literature^(14,31–33,36,37)^.

The dataset also includes information of 81 UFA from 46 patients (42%) and 44 USE from 28 patients. The average number of surgical reconstruction procedures per patient is 1.8 ± 0.6SD, with the nerve transfer (77%) as the most frequent.

Differing from published series, which limit their focus to specific aspects of TBPI, our database provides in the same set a wide range of information concerning this injury. This provides the opportunity of exploring the combination of these descriptors envisioning its deeper understanding.

In conclusion, this is the first report on a public database on TBPI. This is a unique initiative, resulting from a multidisciplinary effort, that stands out by its richness in clinical and epidemiological data and sharing potential. The similarity of this data with other national and international series endorses the quality of the present dataset. TBPI results mainly from traffic accidents, a huge public health problem especially in developing countries, and affects working age population with disabling consequences. In this sense, improving the knowledge on the TBPI can contribute to patient care, support governmental health strategies and give insights into factors related to its prognosis.

## 7 Author Contributions

CBP: questionnaires’ creation, questionnaires’ translation, data collection, data analysis, writing, reviewing

BLR: questionnaires’ creation, questionnaires’ translation, data collection, data analysis, writing, reviewing

JM: questionnaires’ creation, data collection, data analysis

MLR: questionnaires’ creation, questionnaires’ translation, data collection

FFT: questionnaires’ translation, reviewing

LS: questionnaires’ creation, data collection

KRB: intellectual conception, data analysis, writing, reviewing

CDV: intellectual conception, questionnaires’ creation, data analysis, writing, reviewing

## 10 List of abbreviations

INDC-UFRJ: Institute of Neurology Deolindo Couto of the Federal University of Rio de Janeiro
NES: Neuroscience Experiment System
TBPI: traumatic brachial plexus injury
UAA: Unified Admission Assessment
UFA: Unified Follow up Assessment
USE: Unified Surgical Evaluation

## 11 Acknowledgments

We would like to thank José Fernando Guedes Correa, Paulo Leonardo Tavares, José Vicente Martins, Abrahão Baptista, Cissa Nunes Soares, Amanda Nascimento, Carlos Ribas, Cassiano R. N. Santos and Evandro Santos Rocha for their contribution.

## 10 Data Availability Statement

The datasets A Public Database on Traumatic Brachial Plexus Injury for this study can be found in the *Figshare* https://figshare.com/s/6d11e763c9af85b5c95f.

